# SARS-COV-2 induced Diarrhea is inflammatory, Ca^2+^ Dependent and involves activation of calcium activated Cl channels

**DOI:** 10.1101/2021.04.27.441695

**Authors:** Mark Donowitz, Chung-Ming Tse, Karol Dokladny, Manmeet Rawat, Ivy Horwitz, Chunyan Ye, Alison Kell, Ruxian Lin, Sun Lee, Chenxu Guo, Shang Jui Tsai, Andrea Cox, Stephen Gould, Julie In, Steven Bradfute, Nicholas C. Zachos, Olga Kovbasnjuk

**Author notes:** Correspondence to Mark Donowitz, MD, The Johns Hopkins University School of Medicine Ross925, 720 Rutland Avenue, Baltimore, MD 21205, Olga Kovbasnjuk, Ph.D., University of New Mexico Health Sciences Center, IDTC bldg., room 330A, 915 Camino de Salud, Albuquerque, NM 87131.

## Abstract

Diarrhea occurs in 2-50% of cases of COVID-19 (∼8% is average across series). The diarrhea does not appear to account for the disease mortality and its contribution to the morbidity has not been defined, even though it is a component of Long Covid or post-infectious aspects of the disease. Even less is known about the pathophysiologic mechanism of the diarrhea. To begin to understand the pathophysiology of COVID-19 diarrhea, we exposed human enteroid monolayers obtained from five healthy subjects and made from duodenum, jejunum, and proximal colon to live SARS-CoV-2 and virus like particles (VLPs) made from exosomes expressing SARS-CoV-2 structural proteins (Spike, Nucleocapsid, Membrane and Envelope). Results: 1) Live virus was exposed apically for 90 min, then washed out and studied 2 and 5 days later. SARS-Cov-2 was taken up by enteroids and live virus was present in lysates and in the apical>>basolateral media of polarized enteroids 48 h after exposure. This is the first demonstration of basolateral appearance of live virus after apical exposure. High vRNA concentration was detected in cell lysates and in the apical and basolateral media up to 5 days after exposure. 2) Two days after viral exposure, cytokine measurements of media showed significantly increased levels of IL-6, IL-8 and MCP-1. 3) Two days after viral exposure, mRNA levels of ACE2, NHE3 and DRA were reduced but there was no change in mRNA of CFTR. NHE3 protein was also decreased. 4) Live viral studies were mimicked by some studies with VLP exposure for 48 h. VLPs with Spike-D614G bound to the enteroid apical surface and was taken up; this resulted in decreased mRNA levels of ACE2, NHE3, DRA and CFTR. 4) VLP effects were determined on active anion secretion measured with the Ussing chamber/voltage clamp technique. S-D614G acutely exposed to apical surface of human ileal enteroids did not alter the short-circuit current (Isc). However, VLPS-D614G exposure to enteroids that were pretreated for ∼24 h with IL-6 plus IL-8 induced a concentration dependent increase in Isc indicating stimulated anion secretion, that was delayed in onset by ∼8 min. The anion secretion was inhibited by apical exposure to a specific calcium activated Cl channel (CaCC) inhibitor (AO1) but not by a specific CFTR inhibitor (BP027); was inhibited by basolateral exposure to the K channel inhibit clortimazole; and was prevented by pretreatment with the calcium buffer BAPTA-AM. 5) The calcium dependence of the VLP-induced increase in Isc was studied in Caco-2/BBe cells stably expressing the genetically encoded Ca2+ sensor GCaMP6s. 24 h pretreatment with IL-6/IL-8 did not alter intracellular Ca2+. However, in IL-6/IL-8 pretreated cells, VLP S-D614G caused appearance of Ca^2+^waves and an overall increase in intracellular Ca^2+^ with a delay of ∼10 min after VLP addition. We conclude that the diarrhea of COVID-19 appears to an example of a calcium dependent inflammatory diarrhea that involves both acutely stimulated Ca2+ dependent anion secretion (stimulated Isc) that involves CaCC and likely inhibition of neutral NaCl absorption (decreased NHE3 protein and mRNA and decreased DRA mRNA).

## INTRODUCTION

The many recognized clinical manifestations of human COVID-19 infection continue to increase both related to the number of organ systems affected and also to the persistent manifestations that has led to the naming of the prolonged disease, now called post-infectious-COVID-19 or Long Covid. GI manifestations, primarily diarrhea, occur in ∼2%-50% of acute cases (average ∼8%), often early as the presenting complaint but can occur throughout the course of the disease, including during the prolonged phase^1-14^.

GI manifestations are thought to occur from direct luminal rather than systemic contact between the virus and the GI tract. SARS-CoV-2 can be recovered from the lumen of the intestine in spite of being acid sensitive^15,16^ (perhaps related to lack of gastric acid in the period between meals), where it binds and replicates in human enterocytes. Although fecal shedding has been documented in up to 50% cases of infection, this has almost always been looked at as viral RNA rather than live virus, the number of examples of live virus isolation of stool is very limited, and it is unclear if this pool can transmit disease; and this is true during the acute infection as well as during the prolonged phases of COVID-19^8,9,15-17^. Intestinal effects of this virus are not surprising since expression levels of the brush border SARS-CoV-2 receptor, angiotensin-converting enzyme 2 (ACE2), is among the very highest in the human body, being 10-fold higher in intestinal cells than lung cells^16,18,19^. ACE2 is present in the brush border (BB) of enterocytes with the highest concentration in the ileum and much less in the colon and more in the villus than the crypts^18.19^. Intestinal infection by SARS-CoV-2 appears to involve enterocytes with ACE2 not present in goblet cells^14^.

Our understanding of SARS-CoV-2 pathophysiology in intestinal cells involves a dysregulated, aberrant host innate immune response, similar to the response in lung epithelial cells that is characterized by reduced Type I and III interferon (IFN) levels with increased chemokine/cytokine responses^20-25^. There is current uncertainty concerning the role of cytokine storm in the COVID-19 acute respiratory distress syndrome (ARDS) and the role of the inflammatory response in the multiple manifestations of COVID-19. The cytokine response in COVID-19, including the ARDS, is associated with activation of a wide range of pathways, but with lower cytokine levels than in other forms of ARDS. In addition, there is lack of clarity about which cytokines cause symptoms or organ damage in COVID-19, although there is elevation of IL-6 and IL-8 ^26-28, A.Cox, unpublished^.

Important gaps in understanding effects of SARS-Cov-2 on the intestine include lack of adequate clinical descriptions. Unknown characteristics include when in the course of COVID-19 the diarrhea occurs, intestinal site affected, whether the diarrhea is osmotic or secretory, and contribution to morbidity. Equally not understood are mechanistic insights into how the virus causes diarrhea and if and how the inflammatory response that is part of COVID-19 contributes to producing the diarrhea. Also unknown is the role of the GI tract in the many clinical aspects of SARS-CoV-2 infection, including viral replication and disease progression. The current studies make use of a human enteroid model exposed to live virus as well as to two forms of SARS-CoV-2 virus-like particles (VLPs), one of which is competent for ACE2 binding but not fusion, while the other is competent for both receptor binding and membrane fusion/endocytosis. The enteroid model, studied as monolayers, has been used for understanding other host-pathogen interactions and allows the equivalent of luminal pathogen exposure to a model of normal human intestine ^13-15,29,30^.

## RESULTS

### SARS-CoV-2 infects and replicates in human enteroid and colonoid monolayers without compromising the epithelial layer integrity

Initial studies were carried out to confirm human enteroid monolayer expression of ACE2 and viral infection and replication. Differentiated proximal colonic enteroid monolayers expressed ACE2 in the apical surface by IF with expression in colonocytes but not goblet cells. Goblet cells marked by trefoil factor 3 lacked ACE2 (Fig 1A).

**Figure 1.**
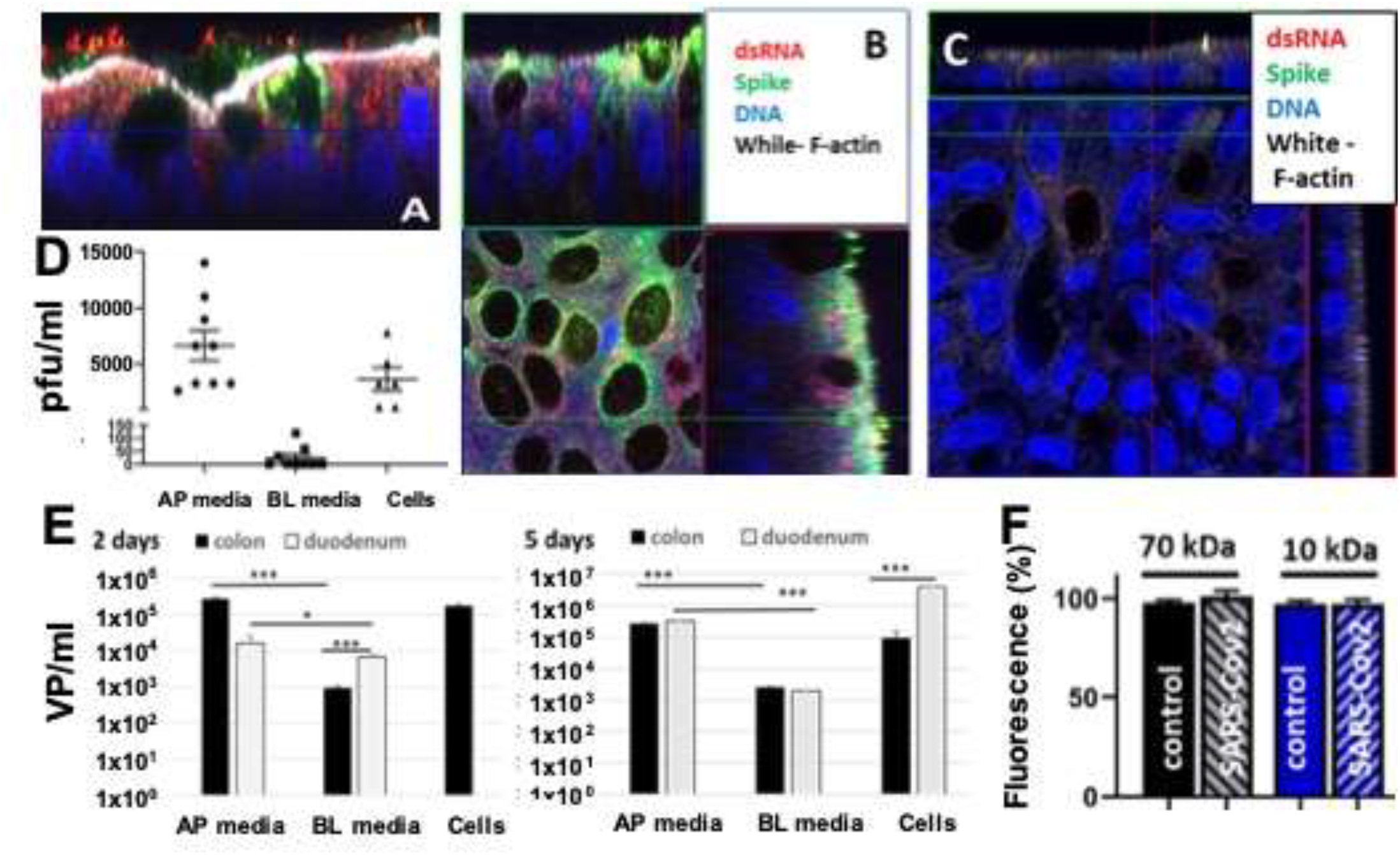
Live SARS-CoV-2 enter and reproduce in human colonoid monolayers. The apical membrane of human small intestinal and proximal colonic enteroid monolayers was exposed to live SARS-CoV-2 (isolate USAWA1/ 2020 SARS-CoV-2; BEI Resources; D Spike protein) for 90 min, washed and then studied for up to 5 days. (A) Colonocytes, but not goblet cells (green = trefoil factor 3) express ACE2 (red). (B) Monolayers were exposed to apical 10^5^ SARS-CoV-2 for 90 min and 48 h later viral entry were confirmed in colonocytes, using double stranded RNA (dsRNA) antibody (red) and viral Spike protein (green). This did not occur in goblet cells 48 h post infection. (C) Spike protein and dsRNA were not detected in uninfected monolayers. (D) Infectious virus was detected in colonoid monolayers as well as in apical and basolateral (BL) media by Vero cell plaque forming assay under conditions in B. Data reported as plaque forming units (pfu). *** p<0.001; n=3. (E) Apical and basolateral media from duodenal or colonoid monolayers apically exposed to live virus (10^6^ PFUs for 90 min) were collected 2 and 5 days post infection. Total RNA was prepared from infected monolayers and expression of viral nucleocapsid was measured by qRT-PCR. The concentration of viral particles (VP) corresponding to RNA levels was calculated based on calibration curves. No VP were detected in mock infected monolayers or media. * p<0.05; *** p <0.001. (F) Viral infection of colonoid monolayers did not affect epithelial permeability of fluorescent 10kDa or 70kDa dextran when studied two days after initial viral exposure. Dextran was added apically and basolateral collection is shown expressed as percent of added dose. Results are mean ± sem. n=3.

Whether human enteroid monolayers could be infected apically with SARS-CoV-2 and whether viral replication occurs in these enterocytes was examined using intestinal segments with lower ACE2 than the ileal expression. Differentiated human proximal colonic and duodenal enteroid monolayers were exposed apically to 10^5^ pfu SARS-CoV-2 in 100 µl (this is the form viral concentrations is presented in this report) for 90 min, then the monolayers were washed and examined at 48 h by IF for presence of Spike protein, viral replication via dsRNA, and presence of live virus in apical and basolateral fluid bathing the enteroids and in enteroid lysates. Shown in Fig 1B, in proximal colonic enteroids (similar results in duodenal enteroids), Spike protein was diffusely present in all enterocytes and particularly in the supranuclear area; while viral dsRNA was present, although in a much more focal distribution, indicating that replication did not occur in all enterocytes in which Spike protein entered. These studies were prolonged for 5 days after a single initial 90 min SARS-CoV-2 exposure (Fig 1B/C). SARS-CoV-2 RNA was present at day 5 in enteroid lysates from duodenum and colon and was also present in the apical and basolateral media. Transepithelial movement of FITC-dextran was measured to determine whether disruption of the enteroid junctional complexes and the accompanying increase in intestinal permeability accounted for presence of basolateral SARS-CoV-2. Studied 48 h after viral apical exposure, there was no change in apical to basolateral movement of 10 and 70 kDa-FITC-dextran compared to untreated control enteroids, indicating no change in intestinal permeability with SARS-CoV-2 exposure. These results support that virus moves from the lumen to the basolateral surface through the epithelial cells, which is the site of the intestinal capillaries and lymphatics. These results are also consistent with the lack of intestinal epithelial pathology detected in 19 COVID-19 patients when examined at the light microscopic level, although studies are limited until now ^*2,7,22,31*^.

### SARS-CoV-2 infection of enteroids causes secretion of proinflammatory cytokines from human enteroid monolayers

COVID-19 is characterized by increased levels of circulating cytokines/chemokines that are responsible for at least some of the severe clinical manifestations. Recent studies have begun to characterize the innate epithelial response of human enteroids to SARS-CoV-2 infection^14-16,20-25^. We evaluated the human enteroid monolayers for changes in SARS-CoV-2 induced cytokine/chemokine production/secretion. SARS-CoV-2 infection of human duodenal enteroids for 90 min induced significant increases in basolateral secretion of cytokines IL-6, IL-8, and the chemokine, MCP-1 (**Fig. 2**) when collected 48 h after viral exposure. Intestinal cells of COVID-19 patients almost certainly have prolonged exposure to increased concentrations of cytokines/chemokines given both this local release and the increased systemic concentrations. To begin to mimic the effect of the prolonged inflammation, we exposed live virus to human enteroid monolayers primed by 24 hour basolateral exposure to the pro-inflammatory cytokines, IL-6 and IL-8. Under these conditions, the epithelial response to SARS-CoV-2 was also significantly increased for TNF-α and IL-1β, further suggesting that the infected intestinal epithelium may contribute to the elevated circulating cytokine levels as well as to the local intestinal environment as part of COVID-19.

**Figure 2.**
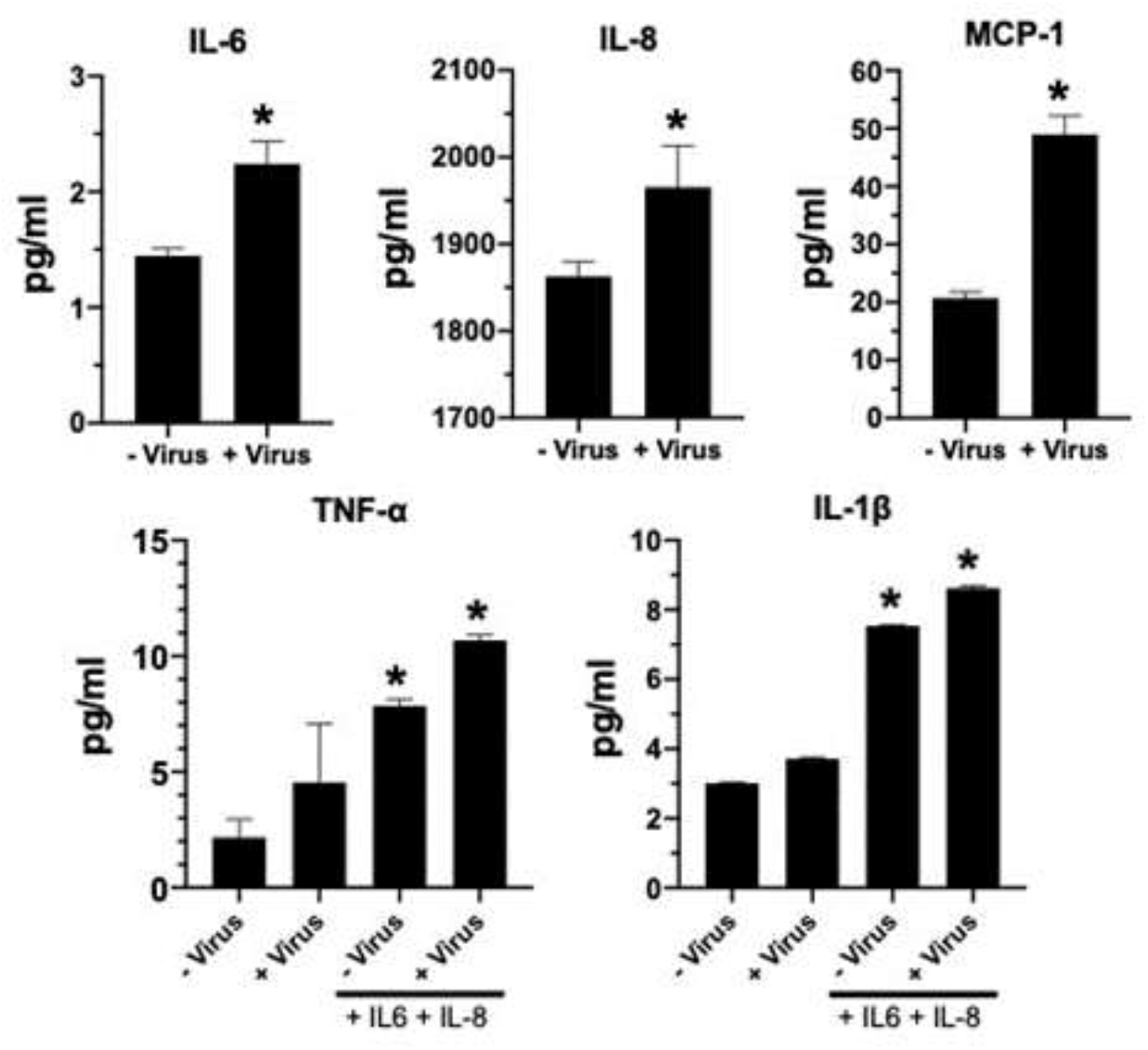
SARS-CoV-2 stimulates epithelial cytokine/ chemokine secretion in human duodenal enteroid monolayers. Differentiated human enteroid monolayers were exposed on the apical surface to SARS-CoV-2 (10^6^) for 90 min, washed out and evaluated 48 h after initial exposure. Studies were performed in the presence or absence of IL-6 and IL-8 (50ng/ml each) for 48 h. Basolateral media were collected and analyzed for secreted cytokines/ chemokines by multiplex ELISA. Results are mean ± sem. *, p<0.05, n= 3.

### SARS-CoV-2-infection of human enterocytes alters expression of ACE2, NHE3 and DRA

Most diarrheal diseases increase stool water by inhibiting the neutral NaCl absorptive process, which is made up of linked neutral exchange proteins, SLC9A3 (the Na/H exchanger NHE3) and SLC26A3 (the Cl/HCO3 exchanger DRA) with or without stimulating the Cl secretory process. To test whether COVID-19 related diarrhea occurs by changes in these transport processes, live SARS-CoV-2 virus was exposed apically to colonoid monolayers for 90 min, and effects on mRNA expression of NHE3, DRA, and CFTR as well as ACE2 were determined using enteroid lysates generated 48 h post-infection. SARS-CoV-2 significantly decreased mRNA levels of NHE3, DRA, as well as ACE2, while CFTR mRNA was unaffected. (Fig. 3A). To determine whether these changes occurred at the protein level, the NHE3 expression pattern was examined in colonoid monolayers by IF/confocal microscopy 48 h after viral exposure (Fig 3B,C). The total and the BB amounts of NHE3 were reduced. While ∼90% of NHE3 under basal conditions is present in the enterocyte apical surface, after SARS-CoV-2 exposure the percent remaining appeared to be more reduced in the apical membrane than in the intracellular pool. These results suggest that at least part of the diarrhea of COVID-19 is due to reduced intestinal NaCl absorption.

**Figure 3.**
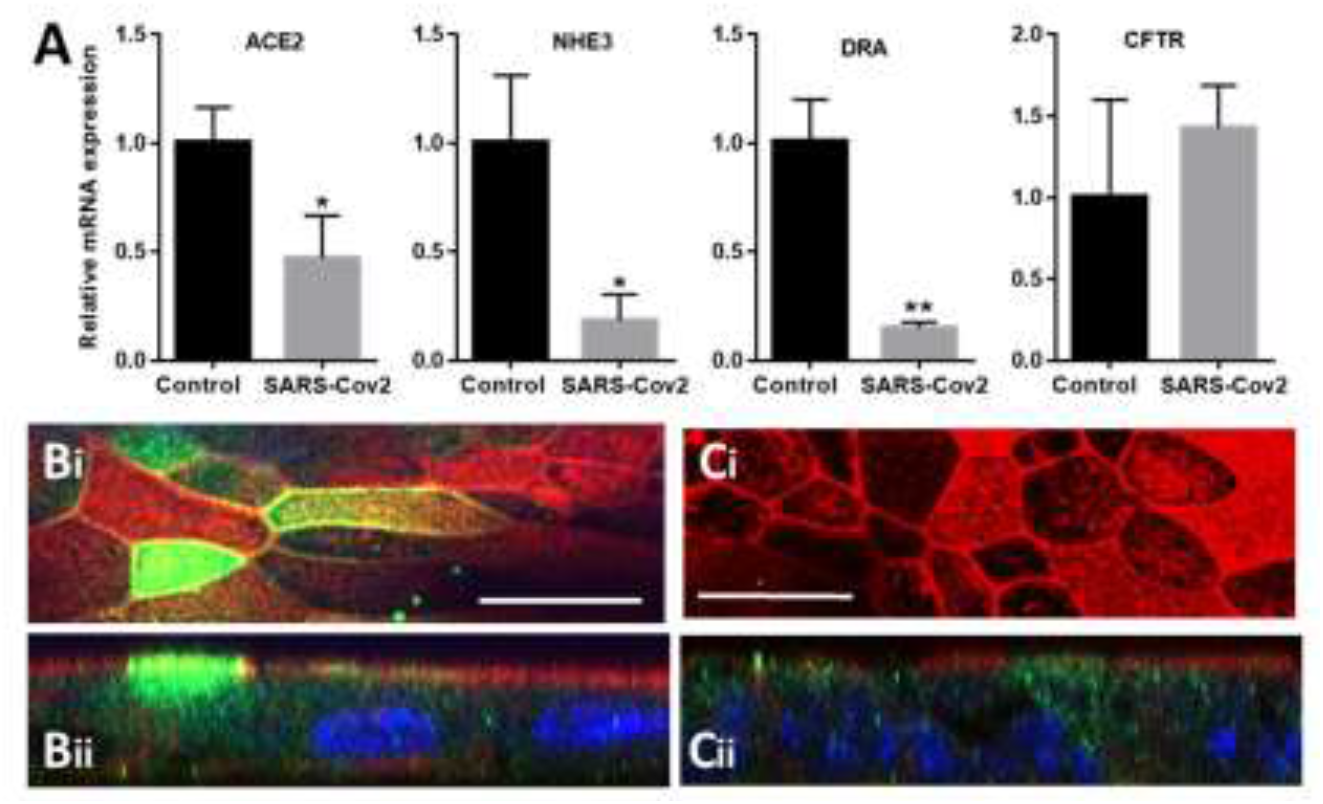
Effect of SARS-CoV-2 infection on the expression of ACE2, NHE3, DRA, and CFTR in human colonoid monolayers. (**A**) SARS-Cov-2 infection for 90 min (10^5^ viral particles exposed apically using enteroid monolayers from 3 different donors) and studied at 48 h reduced mRNA for ACE2, NHE3, and DRA, but not CFTR, Results are means ±sem. *,n=4, p<0.05; (**B**,**C**) Virus alone reduced total NHE3 expression and the percent of total NHE3 in the apical domain. **B**, untreated control. **C**, monolayer infected with 10^5^ virus particles for 90 min and studied 2 days later. Representative confocal XY images at level of the BB (i) and XZ cross sections (ii). NHE3 = green; F-actin = red, nuclei = blue. Scale bar = 20µm. SARS-CoV-2 infection decreased total NHE3 fluorescence intensity measured in projections of at least 100 XZ frames in each of two independent experiments from 8879.1 ±1852.8 in control monolayers to 3092.1 ± 1096.2 (p=0.036); SARS-CoV-2 infection decreased the BB NHE3 fluorescence also measured in at least100 XZ frames from 25410.5 ± 5466.7 in control monolayers to 4823.8 ± 4284.8 (p=0.038) in virus infection monolayers.

### Virus Like Particles (VLPs) that include Spike cause inflammation and reduce transporter expression in human ileal enteroids

SARS-CoV-2 infection involves numerous distinct but overlapping pathologic events. To better understand the different stages of the virus-host interaction, we generated virus-like particles (VLPs) expressing the 4 main structural proteins of SARS-CoV-2: Spike, Nucleocapsid, Membrane, and Envelope in artificial exosomes (small extracellular vesicles). Moreover, two different types of SARS-CoV-2 VLPs were generated, one that expresses the D614G form of Spike present in most current infections, and one that expresses an S-2PP form of Spike that is competent for ACE2-binding but it unable to catalyze fusion between the VLP and cell membranes. As expected, added at time zero and 24 h and sampled at 48 h after initial VLP exposure, S-2PP bound to the enteroid apical surface without evidence of intracellular presence while S-D614G was present both on the surface of the enteroid monolayers and intracellularly (Fig 4A).

**Figure 4.**
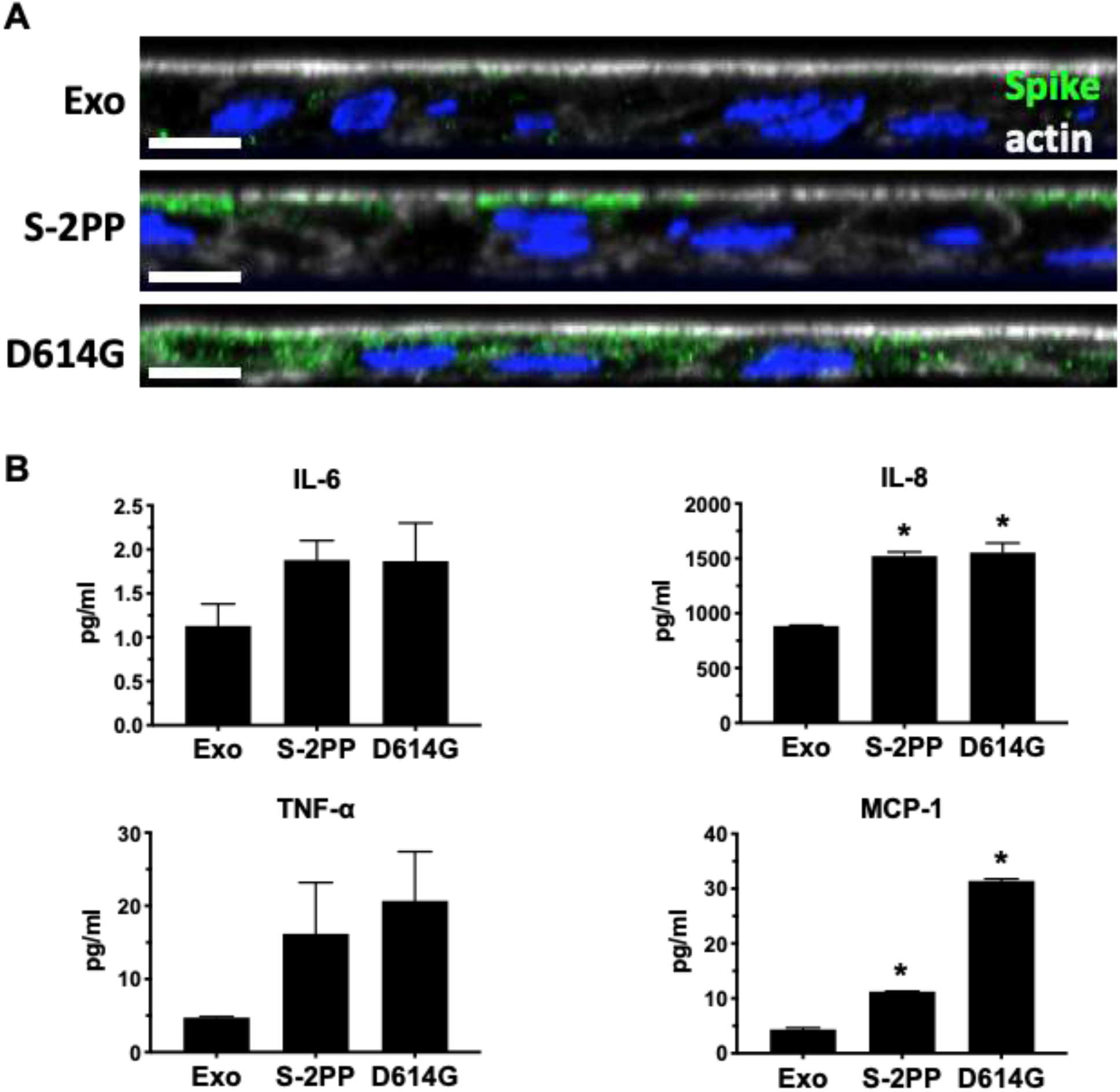
SARS-CoV-2 VLPs directly bind and stimulate epithelial cytokine/chemokine secretion in human enteroid monolayers. (**A**) Immunofluorescence confocal microscopy shows binding of 10^6^ VLP S-2PP to the BB without internalization and both binding and internalization of VLP-G in differentiated human ileal enteroids. VLP were added at times 0 and 24 h and monolayers fixed at 48 h. Exo = empty exosomes; green: = anti-rabbit polyclonal antibody to SARS-CoV-2 Spike protein; white = phalloidin (actin); blue = Hoechst 33342 (nuclei). (B) Differentiated human ileal enteroid monolayers were apically exposed to empty exosomes (Exo); 10^6^ SARS-CoV-2 VLP-S-2PP or VLP-S-G for 48 h, added as in A at times 0 and 24. Basolateral (BL) media were collected and analyzed for secreted cytokines/chemokines by multiplex ELISA. Exposure to VLPs significantly increased secretion of IL-8 and MCP-1, while IL-6 and TNF-α were not significantly increased. *p<0.05. Results are mean ± sem. n=3

Since VLPs designed from other viruses can induce cytokine responses (in the absence of replication)^32^, we hypothesized that direct binding of VLPs to intestinal epithelial cells could induce a cytokine/chemokine response. 48 h after apical exposure of ileal enteroid monolayers at time 0 and 24 h to 10^6^ VLPs, basolateral fluid was analyzed for cytokines. Compared to control exosomes from the same cell line (293F), addition of both forms of SARS-CoV-2 VLPs significantly increased IL-8 and MCP-1 and also increased IL-6 and TNF-α, although the latter two did not reach statistical significance (Fig 4B). Importantly there were no differences between extent of increase caused by Spike binding without entry and Spike binding plus intracellular entry. In addition, VLP effects were determined on the expression of ACE2, NHE3, DRA and CFTR in the same enteroids.

In contrast to effects on cytokine release, there were differences in mRNA expression of ACE2 and transport proteins based on whether VLPs entered the enteroids (Fig 5). Exposure to the fusion incompetent VLPs failed to induce any change in ACE2, DRA or CFTR mRNA and led to only a small decrease in NHE3 message. In contrast, the fusion competent VLPs reduced ACE2, NHE3, DRA and CFTR expression. These were qualitatively similar effects to changes caused by live viral exposure (compare Fig 3A), with additional inhibition of CFTR mRNA. These data suggest that direct binding of VLPs can induce an epithelial cytokine response, which appears to reflect Spike-receptor binding, while fusion competent VLPs lead to additional changes in transport protein and ACE2 expression, even in the absence of viral RNA or viral replication, with all effects mediated by the SARS-CoV-2 structural proteins (S,M,E,N) entering the cell via VLP-cell fusion.

**Figure 5.**
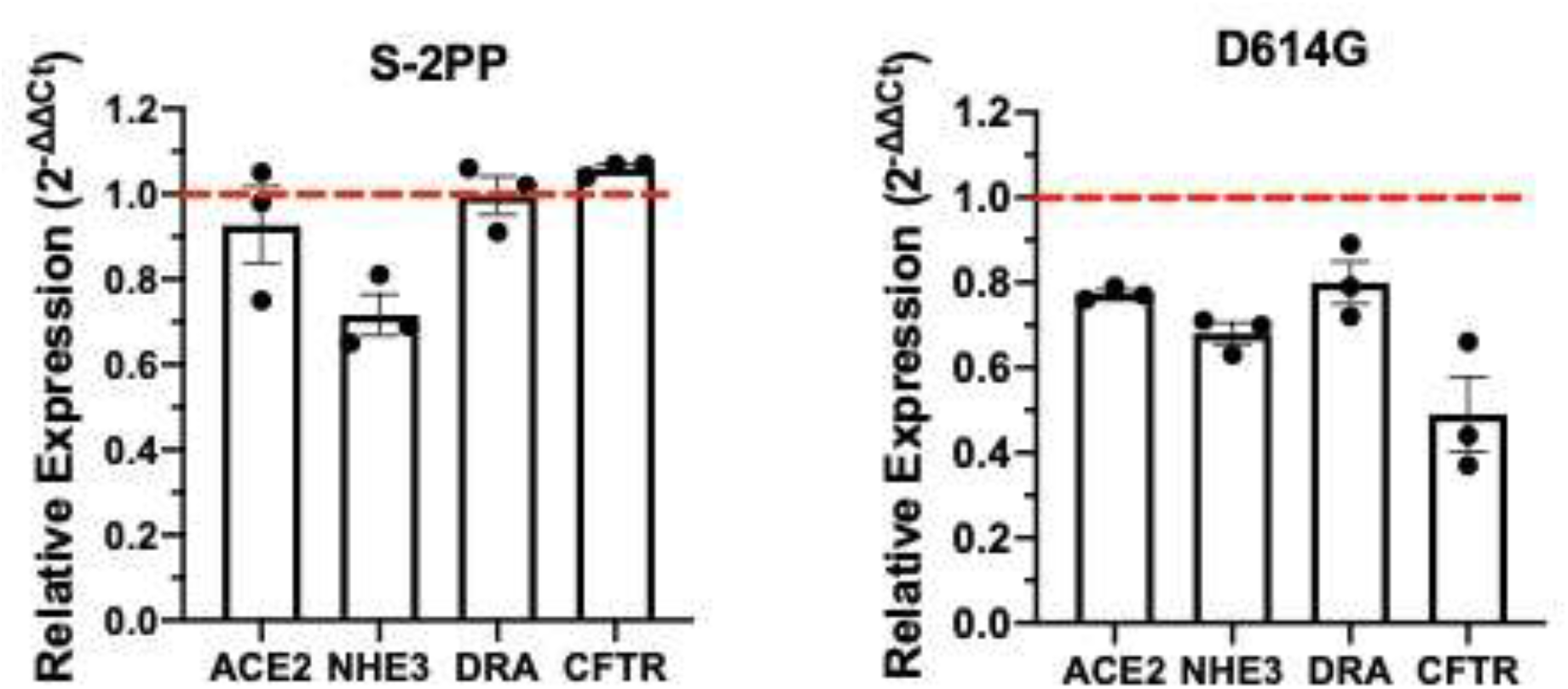
S-12PP and S-D614G forms of SARS-CoV-2 Spike protein differentially affect expression of ACE2, neutral NaCl absorptive transporters and CFTR in human ileal enteroid monolayers. Human ileal enteroid monolayers were exposed apically to VLPs with either S-2PP or S-D614G forms of Spike protein of SARS-CoV-2 in the presence of IL-6/IL-8 (48h, 50 ng/ml). At 48 hours post exposure to VLPs at times 0 and 24 h, total RNA was isolated and mRNA expression of ACE2, NHE3, DRA, and CFTR was measured by qRT-PCR. Red dash = normalization to empty exosome controls. Results are mean ± sem. n=3.

### VLPs cause an inflammatory diarrhea phenotype inducing human ileal enteroid active anion secretion by a Ca^2+^ dependent mechanism that involves CaCC

The VLP-mediated reduction in expression of mRNA for NHE3 and DRA indicates that virus cell binding, fusion, and uptake in the absence of viral replication and production of additional viral proteins may be sufficient to reduce NaCl absorption that is likely to be contributing to the COVID-19 diarrhea. We next tested the hypothesis that SARS-CoV-2 stimulated Cl secretion and that COVID-19 diarrhea was a secretory diarrhea; this hypothesis was in part based on occurrence of diarrhea in some COVID-19 patients who were intubated and not eating. To evaluate VLP effects on active anion secretion, ileal monolayers were studied using the Ussing chamber /voltage clamp technique that measures active electrogenic ion transport, which in diarrheal diseases represents anion secretion, particularly Cl secretion. As shown in Fig 6, studies were performed by addition to the apical surface of VLPs expressing the S-D614G protein. Addition of 10^8^ VLP x3 did not alter Isc (Fig 6A); these enteroids however had rapid increase in Isc with 10 µM forskolin addition (elevates cAMP), indicating the viability of the enteroids. Apical addition of the same concentration of VLPs expressing the S-2PP Spike protein also failed to alter the Isc (Fig 6A). However, given the presence of inflammatory mediators in enteroids exposed to both intact virus and VLP, these studies were repeated with 18-24 h pretreatment with basolateral IL-6 plus IL-8. In the presence of Il-6/IL-8, addition of either VLP apically caused a concentration dependent increase in Isc which occurred both after the third addition of 10^8^ VLPs (data not shown) or a single addition of 3×10^8^ VLP. Shown in Fig 6B-i-v is the D614G Spike protein stimulation of Cl^-^ secretion indicated by the increase in Isc. This was a consistent response with the S-D614G VLP, occurring in 21/23 monolayers tested but was less consistent with the S-2PP VLP, with a change in Isc only occurring n 5/10 monolayers. This IL-6/IL-8 dependent secretion had the following characteristics: a) delayed onset of 7.6 ± 2.7 min (n=5 experiments); b) the increase in Isc was inhibited by the CaCC inhibitor, AO1; c) there was no effect on the Isc response by treatment with the CFTR inhibitor BPO-27; d) the entire Isc response was reversed by basolateral addition of the K^+^ channel inhibitor, clotrimazole. K channels are required to maintain the electrical driving force for Cl secretion and these data support that the increase in Isc represents electrogenic anion secretion; e) the increase in Isc was almost entirely prevented by pretreatment with the intracellular calcium buffer BAPTA-AM (pretreatment 35 mM, 30 min), although after a significant delay some increase in Isc occurred. The last finding is initial evidence of the dependence of VLP-induced anion secretion on an elevation of intracellular Ca^2+^.

**Figure 6.**
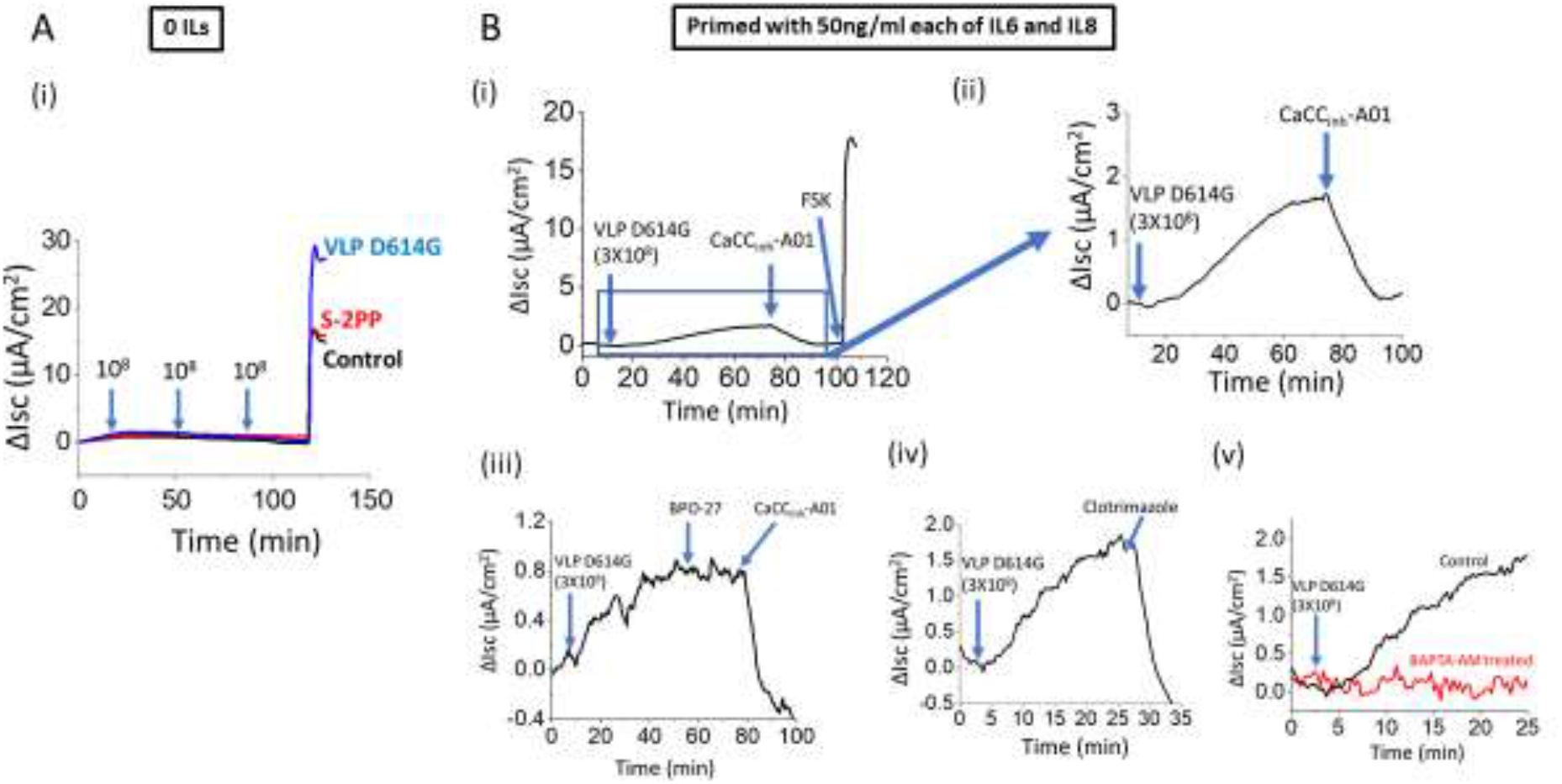
Effect of VLP-S-2PP and VLP S-D614G on Human Ileal Enteroid Monolayer Isc (Anion Secretion). Ussing chamber/voltage clamp studies. (**A**) In absence of IL-6/IL-8, neither VLP altered Isc after three additions of 10^8^. Subsequent addition of forskolin (10μM) gave the predicted increase in Isc. (**B-i-v**) In presence of basolateral IL-6/IL-8 pretreatment (50ng/ml,18-24h), 3 x10^8^ VLP D614G added once increased Isc starting 4-12 min after addition (time to onset of increase in Isc 7.6 ± 2.7 min, n=5 experiments, 9 monolayers). The increase in Isc occurred in 21/23 VLPs. The response to VLP D614G was studied in more detail: **(B-i**,**ii)** The increase in Isc was totally reversed, by the CaCC inhibitor-AO1 (25μM). (**B-iii**) The CFTR inhibitor, BPO-27 (1μM) did not alter the increase in Isc. (**B-iv**)Basolateral exposure to the K channel inhibitor, clotrimazole (30μM) entirely reversed the VLP effect. (**B-v**) Pretreatment with BAPTA-AM (35 μM, 30 min) prevented the VLP-induced increase in Isc.

### VLPs cause an increase in intracellular Ca ^2+^ with a time course similar to that of the VLP stimulation of Cl secretion

To further support a role for intracellular Ca^2+^ in the VLP stimulation of intestinal Cl secretion, the effect of VLP S-D614G was determined on intracellular Ca^2+^ detected by the Ca^2+^ sensor GCaMP6s stably expressed in polarized Caco-2/BBE cells. Using a confocal microscope/environmental chamber to maintain the temperature at 37°C, intracellular Ca^2+^ was determined over the same time course and using the same concentrations of IL-6/IL-8 and apical VLP D614G under which the VLPs altered the Isc in enteroids. Four experimental conditions were compared: VLP effect in the absence and presence of IL-6/IL-8 pretreatment; IL-6/IL-8 treatment alone and time control. Shown in Fig 7, Caco-2 cell Ca^2+^ did not change significantly over the time of the experiment. There was also minimal change in intracellular Ca^2+^ in Caco-2 cells exposed only to IL6/Il-8 overnight (50ng/ml), while a slightly greater increase in Ca2+ occurred with VLP S-D614G treatment alone. In contrast, in Caco-2 cells exposed to IL-6/IL-8 overnight, VLP D614G cause a sustained increase in intracellular Ca2+ that had a delay in time of onset after VLP addition of 9.9 ±1.2 min (n=5 experiments). This is not significantly different than the delay in onset of increased Isc after apical VLP addition. Additionally, intercellular Ca^2+^ waves (ICWs) spreading over multiple cells were detected after VLP S-D614G in the VLP alone and IL-6/IL-8-VLP conditions, suggesting cell to cell signaling. These occurred over a prolonged period, were not seen in the untreated time control or IL-6/IL-8 alone conditions and seems to occur with similar frequency in the VLP and VLP after IL-6/IL-8 condition.

**Fig 7.**
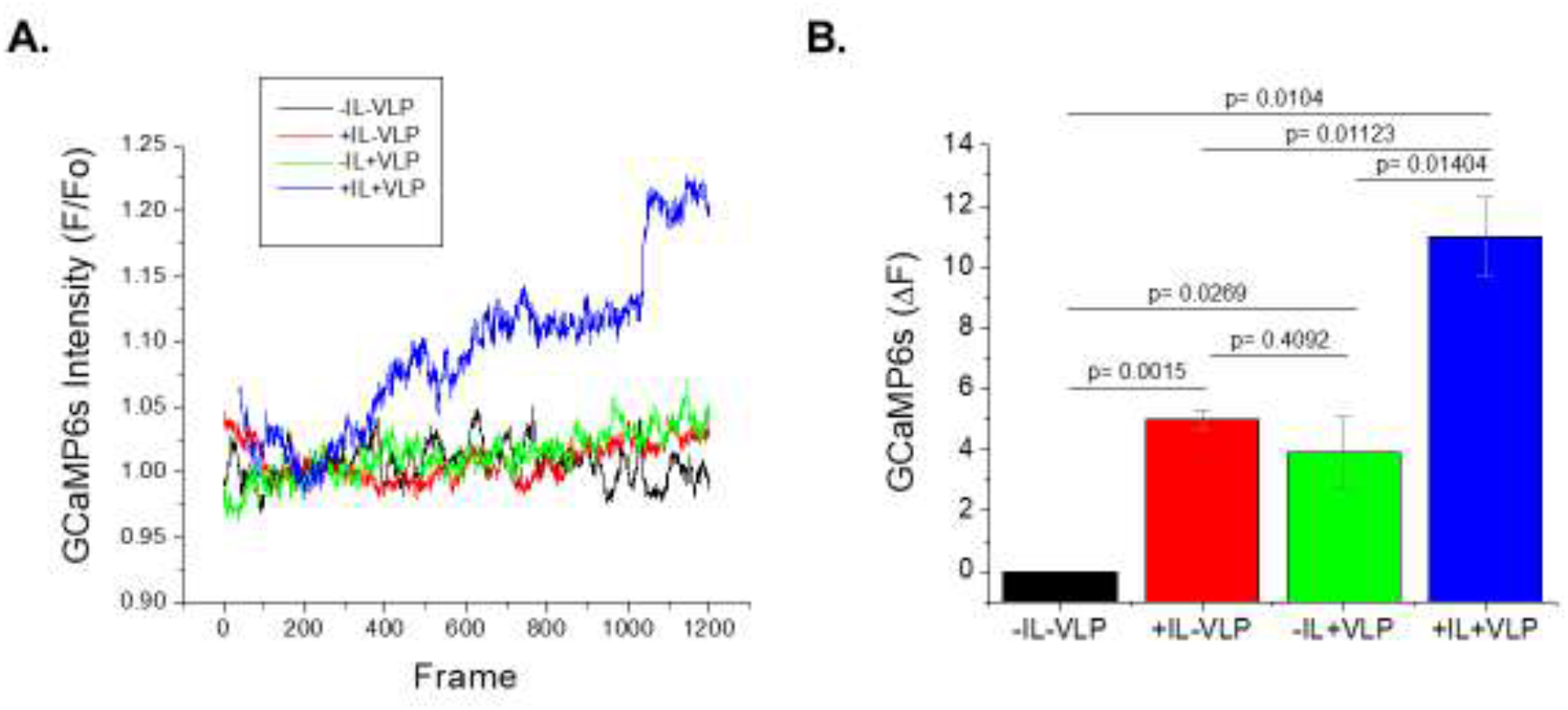
VLP Increases Intracellular Ca ^2+^ in Human Caco-2 Cells but Only with IL-6/IL-8 Exposure. Caco-2/BBE cells stably expressing GCaMP6s were grown on 12 well Transwell inserts for 14 days. Some cells were treated with IL-6 plus IL-8 (both 50ng/ml) for 24h before exposure to VLPs. Live cell images of GCaMP6s fluorescence were determined at 37°C in an Olympus FV3000RS confocal microscope resonant scanner, 520×520 pixels, 16 line averaged (1 second) at 3 second intervals for 60 min without or with VLP addition. Data from full thickness scans were transferred with Metamorph software. A 520 × 520 pixel region of interest was quantified using the Graph Intensities command. Ratios of fluorescence/initial fluorescence is plotted over time for a single experiment in (**A)**. Changes of GCaMP6s intensity (ΔF) from initial baseline prior to VLP addition or similar time in studies without VLP addition were plotted with mean ± sem, n=3 three independent experiments shown in **(B**).

## MATERIALS AND METHODS

### Chemicals, Antibodies and Other Reagents

AO1 and BPO-27 were gifts provided by A. Verkman. Forskolin, clotrimazole, BAPTA-AM (Sigma); IL-6, IL-8 (PepProtech); dextran Alexa Fluor 488 (10kDa) and Alexa Fluor 568 (70kDa) (Invitrogen); anti-dsRNA monoclonal antibodies (clone rJ2; Millipore, cat.# MABE1134); anti-spike rabbit polyclonal antibodies (ProSci, cat # 3525); anti-NHE3 antibodies (Novusbio, Cat# NBP1-82574); anti-ACE2 – monoclonal mouse IgG2A (R&D systems catalog # MAB933); Phalloidin and secondary fluorescent AlexaFluor antibodies from Invitrogen; Vero E6 cells (ATCC® CRL-1586™).

### Enteroid and Colonoid Cultures

Human duodenal and ileal enteroids, and ascending colonic enteroids (colonoids) were derived from intestinal biopsies obtained from normal, healthy donors, as previously described [refs]. Briefly, human enteroids/colonoids were maintained as 3D spheroids in Matrigel in enteroid expansion media containing the growth factors Wnt3a, R-spondin-1, noggin, and EGF necessary for long-term culture [refs]. Enteroid/colonoid monolayers were generated by seeding enteroids/colonoids isolated from Matrigel and fragmented by enzymatic digestion onto 24-well Tranwell inserts (Corning #3470), as described in detail^33^. Results were all from studies on human enteroids or colonoids grown as confluent monolayers on Transwell filters, as we have reported previously^33-39^. Generation and study of human enteroids were approved by the IRBs of JHU SOM (NA_00038329) and Un New Mexico School of Medicine (18-171). Data were obtained from 1 duodenal enteroid line, 2 ileal enteroid lines, and 3 colonic enteroid lines.

### Caco-2/BBE Culture and GCaMP6s Expression

Caco-2/BBe cells were cultured in Dulbecco’s Modified Eagle Medium (DMDM) supplemented with 25 mM NaHCO3, 0.1 mM non essential amino acids, 10% fetal bovine serum, 4 mM glutamine, 100 U/ml penicillin and 100 ug/ml streptomycin in a 5% CO2/95% air atmosphere at 37C Plasmid GP-CMV-GCaMP6s (Addgene plasmid # 40753) was cloned into lentiviral vector pCDH-EF1-MCS-IRES (puro) and lentiviral particles produced. Caco-2 wild-type cells were transduced the the GCaMP6s lentiviral particles and puromycin selected (10 mg/ml). Caco-2 cells stably expressing GCaMP6s were transduced with lentiviral particles produced from lentiviral vector pLV-EF1aa IRES-3HA-CHRM3 (Blasticidin), as described^40^. These Caco-2 cells were maintained under puromycin plus blasticidn containing Caco-2 media. For calcium imaging, these Caco-2 cells were seeded in 12-well Transwell plates (1.2 x10^5^ cells/well) and grown for 12-15 days post confluency; they were serum starved 2-3 h before study.

### VLP-producing cell lines

293F cells were transfected with pS149 and selected in zeocin-containing media, creating the cell line Ftet1, which constitutively expresses the rtTAv16 Tet-on transcription factor from a CMV promoter-driven bicistronic ORF, upstream of a viral peptide and the bleomycin resistance gene. Htet1 cells were converted to SARS-CoV-2 VLP-producing cell lines Ftet1/S226 and Ftet1/CG201 by transfection with Sleeping Beauty transposons that carry 5 separate genes: (1) a puromycin-resistance gene under control of a minimal EF1alpha promoter; (2) SARS-CoV-2 Envelope gene under control of the tetracycline/doxycycline-inducible TRE3G promoter; (3) SARS-CoV-2 Membrane gene under control of the TRE3G promoter; (4) SARS-CoV-2 Nucleocapsid gene under control of the TRE3G promoter; and either (5a) he SARS-CoV-2 Spike-2PP-CSM gene under control of the TRE3G promoter (pS226) or (5b) the SARS-CoV-2 Spike/D614G gene under control of the TRE3G promoter (pCG201) (Spike-2PP-CSM encodes a protein identical to Spike encoded by the Wuhan-1 strain of SARS-CoV-2, with the exception of diproline and cleavage site mutations (986KV987-986PP987 and 682RRAR685-to-682GSAG685); Spike-D614G encodes a protein identical to Spike encoded by the Wuhan-1 strain of SARS-CoV-2 with the exception of the D614G substitution mutation). To generate SARS-CoV-2 VLPs carrying the non-fusogenic yet receptor-binding S-2P-CSM protein, Ftet1/S226 cells were grown in suspension cultures in Freestyle media in the presence of 1 ug/ml doxycycline for 3-4 days. To generate SARS-CoV-2 VLPs carrying the fusogenic, functional Spike-D614G protein, Ftet1/CG201 cells were grown in suspension cultures in Freestyle media in the presence of 1 ug/ml doxycycline for 3-4 days. Starting cell densities were between 0.5-1 × 10^6 cells/ml and the cells were grown in sterile, baffled Erlenmeyer culture flasks at 110 rpm in a humidified, 37°C incubator containing 5% CO_2_.

### VLP purification & characterization

VLPs were purified from Ftet1/S226 and Ftet1/CG201 suspension cell cultures by the following protocol. In brief, cells were removed by centrifugation (5,000 × g for 10 min) and the resulting supernatants were sterilized by gravity filtration through a 220 nm pore diameter filtration, generating clarified tissue culture supernatants. The VLPs were then concentrated ∼300-fold by centrifugal filtration of each CTCS across a 100 kDa cutoff membrane (∼160 mls CTCS to a volume of ∼0.5 ml). The concentrated VLP-containing suspension was then separated by size exclusion chromatography and VLP-containing fractions were collected, pooled, and quantified by nanoparticle tracking analysis using a ParticleMetrix Zetaview camera according to the manufacturer’s instructions. VLP identity was confirmed by immunoblot using antibodies specific for Spike, Membrane, and Envelope.

### Detection of SARS-CoV-2 by RT-qPCR

Presence of SARS-CoV-2 in enteroids and colonoids, as well as in apical and basolateral media were determined using primers and probes (N1, N2 and RP) from the CDC 2019 Real-time Reverse Transcriptase (RT)-PCR diagnostic panel (IDT). Positive controls used in these reactions included SARS-CoV-2 RNA (WA1-USA strain, from UTMB) as well as the 2019-nCoV-N and the Hs-RPP30 control plasmids (IDT). RT-qPCR were performed using TaqPath 1-step RT-qPCR master mix (ThermoFisher) on the ABI StepOnePlus Real-Time PCR system. Reverse transcription and amplification conditions were 25°C for 2 min, 50°C for 15 min, 95°C for 2 min followed by 45 cycles of 95°C for 3 sec and 55°C for 30 sec

### Plaque Assay

SARS-CoV-2 plaque assays in Vero E6 cells were used to assess viral replication. Vero E6 cells were incubated in a 24 well plate until they reached 90% confluency. Supernatant from cultures were mixed with media and added to cells for 2 hours. The supernatant was removed and the cells were overlaid with virus overlay media comprised of equal volumes of a) minimal essential medium/5% FCS/2X antibiotics and b) 2% agarose. After 2 days the overlay was removed, cells were stained with crystal violet, and plaques were counted.

### Enteroid/Colonoid Infection

Confluent enteroid/colonoid monolayers were studied in the undifferentiated state in the presence of conditioned med with Wnt3a, R-spondin1 and noggin and in the differentiated state, 5 days after removal of Wnt, R-spondin1, and noggin, as reported ^38,39^. Enteroids were inoculated apically with either 10^5^ or 10^6^ live SARS-CoV-2 virus (isolate USAWA1/ 2020 SARS-CoV-2; BEI Resources; D Spike protein) for 90 min.; apical media was replaced by virus free media and the infection was monitored for up to 5 days. For studies with VLPs, apical exposure was with 10^6^ or 10^8^ VLPs.

### Cytokine Assays

Cytokine/chemokine levels were assayed using the U-Plex Human Biomarker Group 1 (Cat #: K15067L-1, MesoScale Diagnostics. Wells were washed 3 times with MSD Wash Buffer before incubation with detection antibody solution for 1 hour at room temperature, with shaking. Wells were washed again 3 times before adding 150μl of Read Buffer T to each well. Measurements were obtained on MESO Quickplex SQ 120 Imager using Discover Workbench 4.0 software (MesoScale Diagnostics).

### Ussing Chamber/Voltage Clamp Studies

Ileal monolayers were grown on Transwell inserts and were maintained in the undifferentiated state. These ileal monolayers were switched overnight to CMGF negative medium (Advanced DMEM/F12 medium containing GlutaMAX (2 mmol/L) and HEPES (10 mmol/L)) either with or without 50ng/ml each of IL-6 and IL-8. On the day of experiments, these Transwell inserts were mounted in Ussing chambers (Physiological Instruments, San Diego, CA). The apical and basolateral hemichambers were filled with buffer that was gassed continuously with 95% O2/5% CO2, maintained at 37°C, and connected to a voltage clamp apparatus (Physiological Instruments) via Ag/AgCl electrodes and 3 mol/L KCl agar bridges. Krebs–Ringer bicarbonate buffer consisted of 115 mmol/L NaCl, 25 mmol/L NaHCO_3_, 0.4 mmol/L KH_2_PO_4_, 2.4 mmol/L K_2_HPO_4_, 1.2 mmol/L CaCl_2_, 1.2 mmol/L MgCl_2_, pH 7.4. In addition, buffer in the basolateral hemichamber was supplemented with 4 mmol/L glucose as an energy substrate; buffer in the apical hemichamber was supplemented with 4 mmol/L mannitol to maintain the osmotic balance. To investigate the effect of VLP S-D614G and S-2PP on anion secretion, VLPs were added to the apical hemichamber with a concentration of 10^8^/ml three-times each time with 30min intervals. In subsequent experiments with VLP S-D614G, the VLP particles were added one time with a concentration of 3×10^8^/ml to the apical chamber. Roles of CaCC, CFTR and K channels in VLP D614G stimulated Isc response were studied with a CaCC inhibitor (CaCCinh-A01 20mmol/ml), a CFTR inhibitor (BPO-27, 1µmol/ml) and a K channel blocker (clotrimazole 30mmol/ml), respectively. Effect of BAPTAM-AM (25 uMm, 20 min) pretreated was determined with 25 min preincubation

### Intracellular Ca2+ Measurements withGCaMP6s

Live images of GCaMP fluorescence for intracellular Ca^2+^ measuring were made using the Caco-2/M3 receptor/GcAMP6s monolayers using an Olympus FV3000R confocal microscope (10x objective lens; 2.00x digital zoom; resonant scan with 16 frame average; 0.3 um step size; pinhole 2.3 Airy units, na=0.40) at 37C within a sealed environmental chamber (OkoLab stage top plus transparent shroud incubator) and the associated fluoView (FV31S, v2.1.1.98) acquisition software. Time lapse images were taken approximately once every 3 seconds for 60 minutes. Average fluorescence for the entire frame was generated in MetaMorph Basic (v7.7.3.0) and transferred to Microsoft Excel for normalization before being transferred to Prism 8/9 for graphing and analysis.

### Statistical Analyses

Experiments were generally repeated three times and tested for statistical significance with Student’s paired or unpaired t tests or via ANOVA when three or more comparisons were carried out. p values of <0.05 were consider statistically significant.

## DISCUSSION

While diarrhea is known to occur as a significant clinical component of COVID-19 including acute and long term consequences, little known about the pathophysiology. In fact, studies in animal models, as recently reviewed, and summaries of human disease indicate lack of mechanistic insights^12-14^. The few studies using human enteroids have begun to provide some insights into the pathophysiology and these current studies have confirmed some of the previous enteroid findings and extend mechanistic understanding of the pathophysiology of COVID-19 diarrhea^14-,16^. Importantly, the current studies indicate that the transport effects appear to be the same inhibition of intestinal Na absorption and stimulation of Cl secretion that has been documented to occur in most diarrheal diseases^41-43^ although unique aspects have been revealed. The diarrhea of COVID-19 is the first example of a viral diarrhea that is dependent on the inflammatory response that is part of the disease. The transport processes affected include inhibition of the NaCl absorptive proteins NHE3 and DRA and stimulation of Cl secretion that includes activation of CaCC by a mechanism the is dependent on elevation of intracellular Ca^2+^ which was shown to occur with a time course similar to the time course of the stimulation of active Cl secretion.

The models used in these studies, of human enteroids and colonoids made from healthy human subjects confirmed some previous conclusions using human enteroids. Supporting previous reports, ACE2 expression was present in the apical domain of enteroids made from both small intestine and colon, and was present in both undifferentiated crypt like enteroids as well as differentiated villus like and colon surface cell like epithelial cells but was not present in goblet cells. As reported previously, there was uptake of SARS-CoV-2 and replication in enterocytes measured 48 h after an initial exposure, supporting prolonged viral replication in enterocytes^14,15,41^. As the result of the viral replication, large amounts of viral particles were secreted apically and basolaterally over 5 days of infection^2,7,22,31^. Consistent with previous reports on human intestinal tissue from COVID-19 patients, that intestinal histology is not disrupted by SARS-CoV-2 infection, intestinal permeability measured by transcellular movement of dextran of several sizes was not altered by live viral entry, replication and secretion; this also supports the transcellular movement of SARS-CoV-2 across the epithelium rather than through the paracellular pathway. The apical-to-basolateral surface gradient of secreted virus further support its transcellular secretion route, since increased paracellular movement would lead to similar concentrations on both surfaces. Importantly, live virus was shown to be present in enterocytes 48 h after initial exposure and live virus also present in apical and in one enteroid culture in the basolateral surface, further supporting patient specific viral distribution^31^. The fact that it is live virus that is present, extends previous EM documentation of virus on the basolateral surface, although that study did not require the virus to be capable of further replication ^16^. Since the intestinal mucosal capillaries and lymphatics are present at the base of the epithelial cells, just where live virus (and viral particles) emerges from the basolateral surface of the enterocytes, these results for the first time provide evidence for the possibility that intestinal infection by SARS-CoV-2 is involved in systemic entry of virus via the GI tract, and support that basolateral viral particles (live or not) might contribute to the widespread damage that is part of COVID-19. However, since systemic infection as part of COVID-19 is not felt to be an important part of the pathophysiology of this disease, the clinical relevance of this observation remains undefined.

Further supporting previous studies of COVID-19 pathophysiology, is the increase in cytokine/chemokine release from enterocytes. The inflammatory response in COVID-19, that has now been described in several serial observations over the course of the illness, is complex and involves multiple inflammatory pathways^26-28, A Cox., unpublished.^ Our data suggest that viral replication is not necessary for the production of at least several clinically relevant cytokines with the source being the epithelial cells. Furthermore, we showed that the initial pool of cytokines produced by the epithelial cells are capable of further stimulating viral cytokine production, thus contributing to the accumulating cytokine storm seen in COVID-19. However, we only examined a subset of cytokines shown to be elevated in patients with COVID-19 and our studies were only limited to 48 after initial viral exposure; thus further determinations are needed to define the contributionof intestinal epithelial cells to the inflammatory response seen during the full course of COVID-19.

The changes in transport as part of COVID-19 demonstrated here include 1) reduced expression of NHE3 and DRA which make up neutral NaCl absorption, the major way Na is absorbed from the intestine in the period between meals. The amount of NHE3 was shown to be reduced both at the mRNA and protein level. In addition, the reduced amount of NHE3 seemed to be more of the NHE3 in the brush border compared to the intracellular pool, which suggests that in addition to the reduced total NHE3 amount there was also a change in NHE3 trafficking. NHE3 normally trafficks continually between the brush border and recycling compartment and much of its normal regulation is by protein kinase dependent changes in rates of endo and exocytosis. Further studies will be required to define the role of changes in trafficking of NHE3 and potentially DRA to the diarrhea of COVID-19. 2) stimulated Cl secretion. VLPs induce stimulated Cl secretion by a Ca^2+^ dependent mechanism (prevented by pretreatment with BAPTA-AM), that is delayed with a time course that fits the time for the VLP-induced increase in intracellular Ca^2+^. The VLP effect most consistently required both binding and uptake into the enterocyte, although a less consistent stimulation of Cl secretion occurred with Spike binding alone. This indicates that Cl secretion can occur without the presence of intracellular viral RNA or involvement of other than the viral structural proteins; although whether live virus further induces changes in Cl secretion by the same or different mechanisms than that caused by the VLPs has not been evaluated. One aspect that requires further clarification is the contribution of CFTR. Live virus exposure to the same enteroids used for the VLP studies was not associated with changes in CFTR mRNA, while S-D614G VLP caused a decrease; since the VLP stimulation of Cl secretion did not involve CFTR, similar studies with live virus are required to determine if CFTR contributes to SARS-CoV-2 induced Cl secretion. It is worth noting, however, that while CFTR is the major contributor to secretory diarrheas like cholera, it appears to play less of a role or not to be stimulated in inflammatory diarrheas^42-44^, which appears to characterize the diarrhea induced by SARS-CoV-2. While these results implicate a calcium activated chloride channel (CaCC) in COVID-19 diarrhea, we have not defined which isoform is involved; transport has only been characterized in ileum, and it is likely that different CaCCs are involved in anion secretion in different intestinal segments. The mechanism of the demonstrated elevated intracellular Ca2+ has not been clarified and represents an important area for further study.

The mechanism of COVID-19 diarrhea appears to require the molecular interplay among inflammatory cytokines, the viral receptor, and several apical and basolateral ion transport proteins as well as activation of intracellular Ca2+ signaling (Fig 8). The changes in Na and Cl transporters with SARS-CoV-2 required the related inflammatory environment, modeled by exposure to IL-6/IL-8. This was necessary but not sufficient to induce the changes. Neither IL-6 nor IL-8 elevate intracellular Ca^2+,^ based on multiple previous as well as the current measurements, although one or both along with elevated Ca2+ were necessary for the VLP stimulated Cl secretion. The role of the COVID-19 inflammation and elevated intracellular Ca2+ is less clear for NHE3 regulation and additional studies will be required to define the role of the complex inflammatory state of COVID-19 and the changes that occur during the multiple stages of the disease on Na and Cl transport. Moreover, the transport protein studies have not included all the transporters potentially affected in COVID-19. For instance ACE2 interacts with a brush border amino acid tansporter, SLC6A19 (BoaTi)^45^; and both short and long term changes in ACE2 in COVID-19 could alter amino acid absorption, with potential nutritional implications, especially for the prolonged aspects of this disease. In addition, even though characterization of the changes in Na and Cl transport which occur throughout COVID-19 are incomplete, the results obtained already suggest potential drug targets, while mechanistic insights may be relevant to SARS-CoV-2 effects on other epithelial cells.

**Fig 8.**
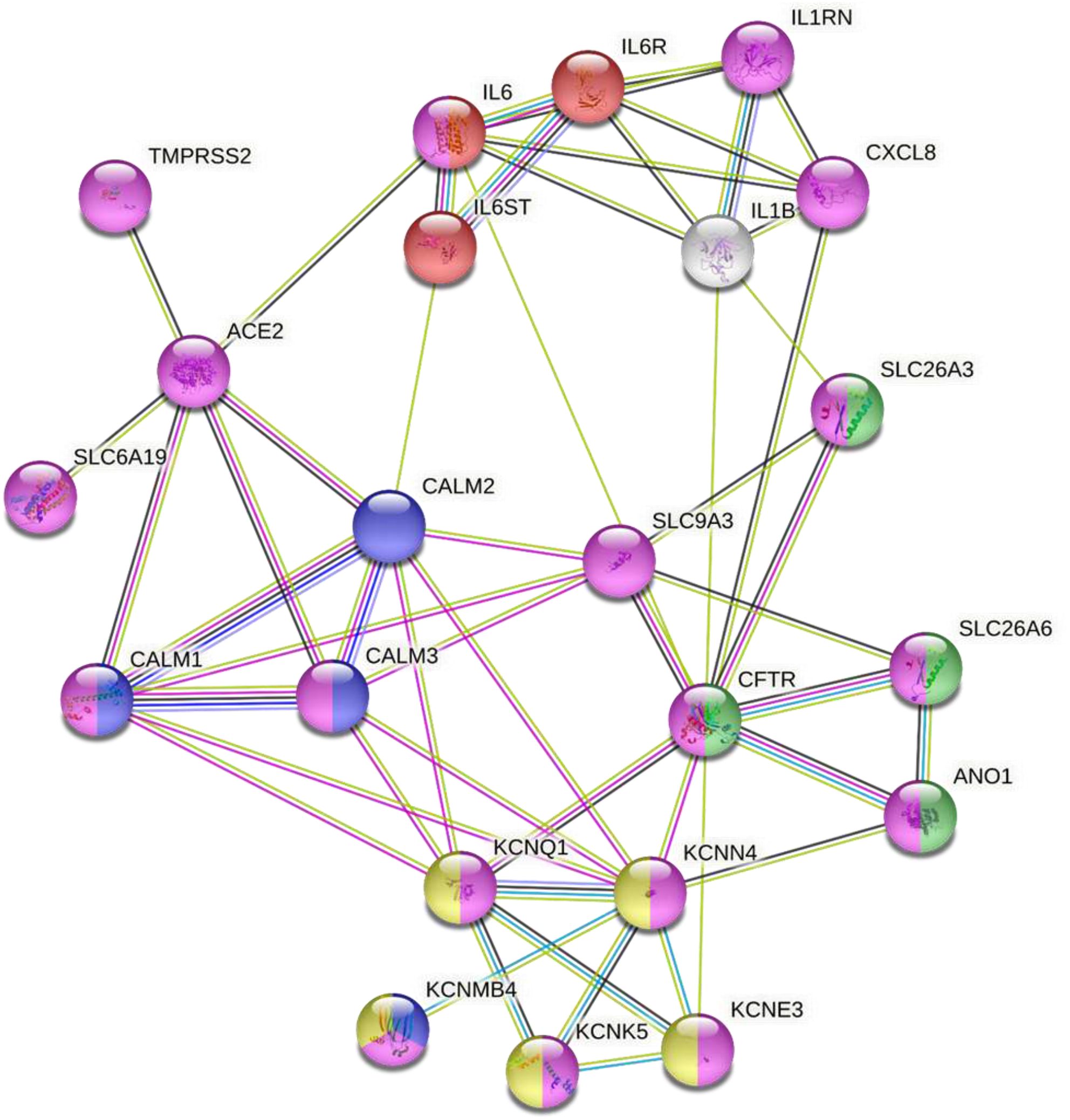
Modeling the molecular mechanisms of SARS-Cov-2 induced diarrhea using a functional interactome created from a small number of proteins that are known to be modulated by the viral infection of human intestinal epithelium (https://string-db.org/). This analysis is based on the most inclusive gene ontology transport pathway (GO:0006810; false discovery rate (FDR) = 2.30e-07); magenta) and includes 21 functionally and physically interacting proteins that are potentially involved in virus-induced signaling that triggers the Isc response in human intestinal epithelium. This model indicates that NHE3 plays a central role in signal transduction between GO:0070102 - the interleukin-6-mediated signaling pathway (red; false discovery rate (FDR)= 4.08e-0) and GO:1902476 - chloride secretion (green, FDR = 5.96e-0). This cross-talk between interleukins and Cl secretion might be mediated by PAT-1 in a CFTR-independent or -dependent manner, leading to ANO1 activation. NHE3 is also crucial for the signal transduction from virus receptor ACE2 via intracellular calcium stores to a membrane Cl/bicarbonate transporter and basolateral voltage-gated potassium channel complex (yellow - CL:8927; FDR = 5.91e-07). The IL-6 functional network (red) together with IL-8 (CXCL8) influences the functions of the SARS-Cov-2 receptor complex and membrane transporters (magenta) which cause intestinal anion secretion via CaCC and via the Ca2+ signaling cascade (blue) that affects the activity of Ca^**2+**^ - or cAMP-mediated basolateral K+ channels (yellow). Following string colors that connect the proteins represent the interactions based on: turquoise - from curated databases; magenta – experimentally determined; green – gene neighborhood; blue – gene co-occurrence; light green – text mining; black – co-expression; light purple – protein homology.

To better understand the relationship between viral infection and possible pathophysiologic mechanisms of diarrhea, we used the STRING functional protein association network to model the potential physical and/or functional molecular links between virus receptor ACE2, proinflammatory cytokines elevated in the intestinal epithelium due to viral infection, and the virus-modulated ion transporters as well as their interacting partners (Figure 8). Our model suggest that IL-6 functionally interacts with ACE2 and CFTR and these interactions affect NHE3 and DRA, and to a less extent, SLC6A19. IL-8 (CXCL8) might affect the ion transporters similarly to IL-6, through direct interactions with IL-6 and CFTR. This model also suggests that ACE2 binding to calmodulin^46^ and calmodulin association with NHE3 might provide insights into at least part of the Ca^2+^ signaling involvement in COVID-19 diarrhea. While a role for CFTR in the COVID-19 diarrhea has not been identified, the current studies have shown that Cl secretion occurs through one or more CaCCs. Shown in Fig 8, ACE2 through the interactions with NHE3 is predicted to transduce the IL-6 signaling to SLC26A6 (PAT-1), a small intestinal apical Cl-/bicarbonate exchanger present in the small intestine particularly in the proximal small intestine, which potentially could take part in activation of ANO1. This bioinformatics based model requires testing and PAT-1 plays a minor role in distal small intestinal transposrt in which the VLP studies of ion transport were performed. This model suggests that both CFTR and CaCCs might contribute to SARS-CoV-2-induced Cl-secretion and intracellular Ca2+ flux could be modulated by ACE2 signaling. Additionally, several basolateral K-channels are predicted to be involved in this viral diarrhea. We suggest that this model might be of use as a roadmap to further experimentally examine the disruption of intestinal transport that contributes to the pathophysiology of COVID-19 diarrhea.

## Acknowledgements

This study is partly supported by National Institutes of Health grants R01-DK-26523, R01-DK-116352, R24-DK-64388, and P30-DK-89502. The authors thank George McNamara (Johns Hopkins University) for assistance in confocal imaging.

